# Investigating the Lipid Selectivity of Membrane Proteins in Heterogeneous Nanodiscs

**DOI:** 10.1101/2021.08.17.456692

**Authors:** James E. Keener, Hiruni S. Jayasekera, Michael T. Marty

## Abstract

The structure and function of membrane proteins can be significantly impacted by the surrounding lipid environment, but membrane protein-lipid interactions in lipid bilayers are often difficult to study due to their transient and polydisperse nature. Here, we used two native mass spectrometry (MS) approaches to investigate how the *Escherichia coli* ammonium transporter (AmtB) and aquaporin Z (AqpZ) selectively remodel their local lipid environment in heterogeneous lipoprotein nanodiscs. First, we used gas-phase ejection to isolate the membrane protein with bound lipids from heterogeneous nanodiscs with different combinations of lipids. Second, we used solution-phase detergent extraction as an orthogonal approach to study membrane protein remodeling of lipids in the nanodisc with native MS. Our results showed that Triton X-100 and LDAO retain lipid selectivity that agrees with gas-phase ejection, but C8E4 distorts some preferential lipid interactions. Both approaches reveal that AmtB has a few selective binding sites for phosphatidylcholine (PC) lipids, is selective for binding phosphatidylglycerols (PG) overall, and is nonselective for phosphatidylethanolamines (PE). In contrast, AqpZ prefers either PC or PG over PE and prefers PC over PG. Overall, these experiments provide a detailed picture of how membrane proteins bind different lipid head groups in the context of mixed lipid bilayers.

## INTRODUCTION

Membrane proteins play crucial roles in cellular processes and represent the majority of drug targets.^1–3^ Membrane protein structure and function can be heavily influenced by interactions with the lipid environment, either by global bilayer properties or through direct lipid interactions.^4–5^ This interplay between proteins and lipids drives membrane organization and structure. For example, membrane proteins can be recruited to membrane microdomains based on local bilayer properties like curvature or thickness.^6–7^ Conversely, membrane proteins can remodel their local lipid environment by selectively binding different lipid types.^8–9^

Molecular dynamics simulations have provided significant insights about lipid remodeling,^10^ but membrane protein-lipid interactions remain challenging to study experimentally due to their dynamic and heterogeneous nature.^11^ Cryo-EM and X-ray crystallography provide high-resolution snapshots of stable membrane protein-lipid complexes with non-annular lipids. However, these structural methods are less suitable for annular lipids that dynamically exchange with bulk lipids in the surrounding bilayer.

Native mass spectrometry (MS) has been used to quantify, distinguish, and identify different kinds of bound lipids surrounding membrane proteins,^12–16^ but these experiments are generally limited to probing a small number of lipids in detergent micelles. To investigate a wider range of lipid interactions, we previously used nanodiscs, discoidal lipid bilayers encircled by two membrane scaffold protein (MSP) belts, to study membrane proteins in lipid bilayers with a larger number of bound lipids. However, past studies have only explored a single lipid or pair of lipids.^17–18^

Here, our goal was to investigate how membrane proteins remodel their surrounding lipid bilayer in more complex mixed lipid environments. We chose the ammonium transporter trimer (AmtB) from *Escherichia coli* (*E. coli*) because lipid interactions with this protein affect its stability and function. Specifically, AmtB binds and is stabilized by phosphatidylglycerol (PG) lipids.^8^ Furthermore, molecular dynamics simulations and activity assays revealed that AmtB requires PG lipids for proper function.^19^ To further validate the method, we also tested *E. coli* aquaporin Z (AqpZ).

We performed native MS of the membrane protein-lipid complexes following gas-phase ejection or solution-phase detergent extraction of the protein from mixed lipid nanodiscs composed of binary and ternary mixtures of palmitoyl-oleoyl-phosphatidylethanolamine (POPE), palmitoyl-oleoyl-phosphatidylcholine (POPC), and palmitoyl-oleoyl-phosphatidylglycerol (POPG). Ejection of AmtB revealed an overall selectivity in average bound lipid composition for POPG, a few tightly retained POPC lipids, and no selectivity of POPE. Detergent extraction using Triton X-100 showed similar selectivity trends, but C8E4 extraction showed little to no lipid selectivity, illustrating that detergents can affect membrane protein-lipid interactions when extracting from lipid bilayers. AqpZ showed similar detergent effects but an overall affinity for either POPC or POPG.

## METHODS

### Protein Expression and Purification

HIS-MBP-TEV-AmtB, AqpZ-TEV-GFP-HIS, and MSP1E3D1 were expressed in *E. coli* and purified as previously described.^17, 20^ Briefly, AmtB and AqpZ were detergent extracted overnight with *n*-dodecyl-β-d-maltopyranoside (DDM, Anatrace) and *n*-octyl-β-d-glucopyranoside (OG, Anatrace) following isolation of membranes by ultracentrifugation. They were then purified by immobilized affinity chromatography (IMAC), followed by size exclusion chromatography (SEC) on a HiLoad 16/600 Superdex 200 pg (Cytiva) with 0.025% DDM. MSP1E3D1 was purified by IMAC using established protocols.^21^

### Nanodisc Assembly and Purification

AmtB and AqpZ nanodiscs were assembled using MSP1E3D1(–), with the polyhistidine tag removed by TEV protease, and purified as previously described.^17–18, 20^ 1-Palmitoyl-2-oleoyl-*sn*-glycero-3-phosphocholine (POPC), 1-palmitoyl-2-oleoyl-*sn*-glycero-3-phospho-(1′-rac-glycerol) (POPG), and 1-palmitoyl-2-oleoyl-*sn*-glycero-3-phosphoethanolamine (POPE) lipids from Avanti Polar Lipids were dissolved in chloroform and quantified by phosphate analysis. Lipids in chloroform were mixed to a molar ratio of 1:1 or 1:1:1, dried overnight, and resuspended in 100 mM sodium cholate (Sigma Aldrich) to a final lipid concentration of 50 mM. Lipids, MSP, cholate, and AmtB/AqpZ were mixed and incubated on ice for approximately 1 hour. Following addition of Amberlite XAD-2 beads (Sigma Aldrich), the reconstitution mixture was incubated at 4 °C overnight on an orbital shaker. Nanodiscs were then purified by IMAC followed by SEC using a Superose 6 10/300 Increase GL (Cytiva). Nanodiscs were incubated with TEV protease at 4 °C overnight to cleave the HIS-MBP or GFP-HIS tag. Nanodiscs were purified by another round of IMAC and SEC, and peak fractions were concentrated to 1–5 μM nanodisc. Samples were either analyzed immediately or flash frozen at −80 °C for storage. Error bars correspond to three replicate nanodisc assemblies that were prepared separately for each lipid composition.

### Native MS of Nanodiscs

For ejection-based native MS, nanodiscs were gently mixed 19:1 v/v with neat glycerol carbonate at >90% purity (Tokyo Chemical Industry Co., Inc.).^17–18^ The mixture was incubated for several minutes at room temperature prior to native MS.

For detergent extraction, stock C8E4 (Anatrace), lauryldimethylamine oxide (LDAO, Anatrace), and Triton X-100 (Sigma Aldrich) solutions were prepared by diluting to 20× the critical micelle concentration (CMC) in water. Nanodiscs were gently mixed 9:1 v/v with stock detergent solutions to yield a final detergent concentration of 2× CMC, incubated on ice for 3 minutes, and then analyzed by native MS.

Compared to gas-phase ejection (Figures S-1–S-4), solution-phase detergent extraction yielded AmtB trimer that were more stable with less subunit dissociation (Figures S-5–S-10). AmtB trimer dissociation was minimal for Triton X-100 and absent for C8E4. Similar trends were observed for AqpZ (Figures S-11–S-13), which showed minimal dissociation with detergent extraction. We also observed that Triton X-100 preserved a larger number of lipids bound to AmtB, demonstrating that different detergents can be used to isolate different numbers of bound lipids. LDAO retained fewer lipids bound to AqpZ than C8E4.

Native MS of nanodiscs was performed as previously described^17^ using a Q-Exactive HF Orbitrap mass spectrometer (Thermo Fisher Scientific, Bremen) equipped with Ultra-High Mass Range modifications.^22^ Nano-electrospray ionization was performed in positive ion mode using borosilicate needles pulled with a P-1000 micropipette puller (Sutter Instrument, Novato, CA). AmtB nanodiscs were analyzed with a range of 2,000–30,000 *m/z* at a resolution setting of 15,000. Collision voltage was applied in the HCD cell and increased from 0 to 200 V in 20 V increments at 1- or 2-minute acquisitions for each step. Important instrument settings included: 1.1–1.5 kV capillary voltage, 200 °C capillary temperature, and a trapping gas pressure of 7. Nanodiscs with Triton X-100 were analyzed with a trapping gas pressure of 10 when the membrane scaffold protein (MSP) signal was relatively high. Ternary nanodiscs with glycerol carbonate were analyzed with a capillary temperature of 300 °C to aid with desolvation. Moreover, ternary and 50% POPG:POPE nanodiscs with glycerol carbonate were analyzed with 0–50 V of source fragmentation to aid with desolvation. Data for AmtB ejected from 50% POPC:POPG nanodiscs was collected and described previously.^18^ AqpZ nanodiscs were analyzed with a range of 2,000–25,000 *m/z* and 0-50 V source fragmentation but otherwise identical instrument settings to AmtB.

### Native MS Data Analysis

Analysis of native MS data was performed as previously described^18^ with slight modifications. MS data was analyzed using both UniDec^23^ and MetaUniDec.^24^ Important deconvolution parameters were as follows: mass range of 29–200 kDa for AmtB nanodiscs and 20–200 kDa for AqpZ nanodiscs, charge range of 1–25, mass sampled every 1 Da, and a peak full width at half maximum of 2.5 Th using a Gaussian peak shape function. The charge, point, and mass smooth widths were all set to 1. Mass smoothing was used with mass differences corresponding to average lipid masses of 754.5 Da, 733.5 Da, 739 Da, and 742.4 Da, for POPC:POPG, POPG:POPE, POPC:POPE, and POPG:POPC:POPE nanodiscs, respectively. For reference, the masses of the pure lipids are POPC 760 Da, POPG 749 Da, and POPE 718 Da. Prior native MS studies with heterogeneous intact empty nanodiscs have shown that using two lipids with similar masses limits the complexity of spectra because nanodiscs with different combinations of lipids produce a clean series of overlapping peaks, where the average lipid mass can be used to determine lipid composition. ^25–27^

Lipids bound to proteins can undergo preferential gasphase dissociation at high levels of activation that is related to the gas-phase basicities of the headgroups.^28^ To avoid these and other gas-phase artifacts, we chose voltage ranges with lower levels of activation. The lowest voltage for each lipid mixture was set to where the peaks were first well-resolved. The maximum voltage for each lipid mixture was set to capture low numbers of bound lipids without observing significant artifacts or dissociation of the protein complex. For ejection-based native MS of AmtB nanodiscs, the voltage ranges for POPC:POPE, POPG:POPE, POPC:POPG, and POPG:POPC:POPE were 60–160 V, 0–160 V, 60–200 V, and 20–180 V, respectively. For detergent extraction, the voltage ranges for POPC:POPE, POPG:POPE, POPC:POPG, and POPG:POPC:POPE were 100–160 V, 100–160 V, 100–200 V, and 100–180 V, respectively. At these levels of activation, less than 10% of the signal was for dimer and monomer. We did not observe lipid enrichment in line with the predicted gas-phase basicity of headgroups at these voltages, suggesting that we have avoided artifacts.

For ejection-based native MS of AqpZ nanodiscs, the voltage ranges for POPC:POPE, POPG:POPE, and POPC:POPG were 120–160 V, 120–160 V, and 140–180 V, respectively. For detergent extraction with LDAO, the voltage ranges for POPC:POPE, POPG:POPE, and POPC:POPG were 120–180 V, 140–200 V, and 140–200 V, respectively. For detergent extraction with C8E4, the voltage ranges for POPC:POPE, POPG:POPE, and POPC:POPG were 60–120 V, 60–160 V, and 60–160 V, respectively. We observed slightly more subunit dissociation during ejection of AqpZ compared to AmtB, but it was still generally less than 30% of the signal.

Spectra collected over these voltage ranges were deconvolved to zero-charge mass spectra and averaged to provide a comprehensive span of different membrane protein-lipid complexes. We extracted the center of mass for each peak from the deconvolved mass distribution corresponding to a membrane protein-lipid complex. Only intensities above 50% of the maximum around each peak were used for extraction to minimize distortions from the baseline or noise. Following subtraction of the membrane protein mass, we divided by the number of bound lipids to calculate the average lipid mass for a given number of bound lipids.

We observed small differences in the mass of the membrane protein, likely due to adduction or errors in mass accuracy, and corrected for this using a linear regression of number of bound lipids versus the center of mass for the peaks. Similar to our previous study,^18^ the correction was typically less than the mass of one water or ammonium molecule. Additionally, the average masses of bound lipids are not reported for 1 and 2 bound lipids, because average mass uncertainties are larger at lower numbers of bound lipids. Different peaks were present at different collision voltages, so the overall average mass for each peak was calculated by averaging each collision voltage step weighted by the squared intensity at that voltage. The purpose of squaring the signal was to focus more on high-intensity signals relative to low-intensity signals that can be influenced by noise and baseline. The number of lipids bound to the membrane protein was somewhat different between lipid compositions and between gas-phase and solution-phase extraction methods. To facilitate more direct comparison, the average lipid masses were plotted against the minimum common number of lipids observed for either binary or ternary mixtures. Because the lipids are close in mass, the number of bound lipids for each peak can be counted unambiguously.

To demonstrate that the average lipid mass accurately reports the bound lipid composition, we simulated mass spectra corresponding to AmtB-lipid complexes with different ratios of bound POPC:POPE (Figure S-14), including a spectrum with a bimodal distribution of ratios. We assumed a binomial distribution of the two bound lipid species. The simulated spectra were analyzed with the same method described above to determine the average lipid mass and then bound lipid composition. Strong agreement of the simulations with known values demonstrates the robustness of the data analysis approach.

## RESULTS AND DISCUSSION

### Ejection of AmtB from Two-Component Lipid Nanodiscs Reveals Lipid Enrichment

To investigate how AmtB selectively remodels its surrounding lipid environment, we used native MS to eject AmtB-lipid complexes from nanodiscs with binary mixtures of POPC, POPG, and POPE. POPC is not present in *E. coli* membranes but was included because it is frequently used in membrane mimetics. For brevity, we refer to each lipid by its headgroup abbreviation: PC, PG, and PE. AmtB was assembled into nanodiscs with 50% PC:PE, 50% PG:PE, and 50% PC:PG. The nanodiscs were analyzed by native MS using glycerol carbonate as a supercharging reagent to facilitate ejection of AmtB from the nanodisc.^17^ Collisional activation was gradually applied to eject AmtB with bound lipids (Figure 1A), and data were deconvolved to zero-charge mass spectra. Representative raw (Figure S-1 and S-2) and zero-charge mass spectra (Figures 1B, 1C, 1D) are shown for each lipid mixture where each peak corresponds to AmtB with a specific number of bound lipids. We measured the average mass of bound lipids for each peak by subtracting the mass of AmtB and dividing the difference by the number of bound lipids. If there were no enrichment, the average lipid mass would be equal to the average of the two lipids. Any shifts in the average lipid masses indicate preferential binding of either the lighter or heavier lipid.

**Figure 1.**
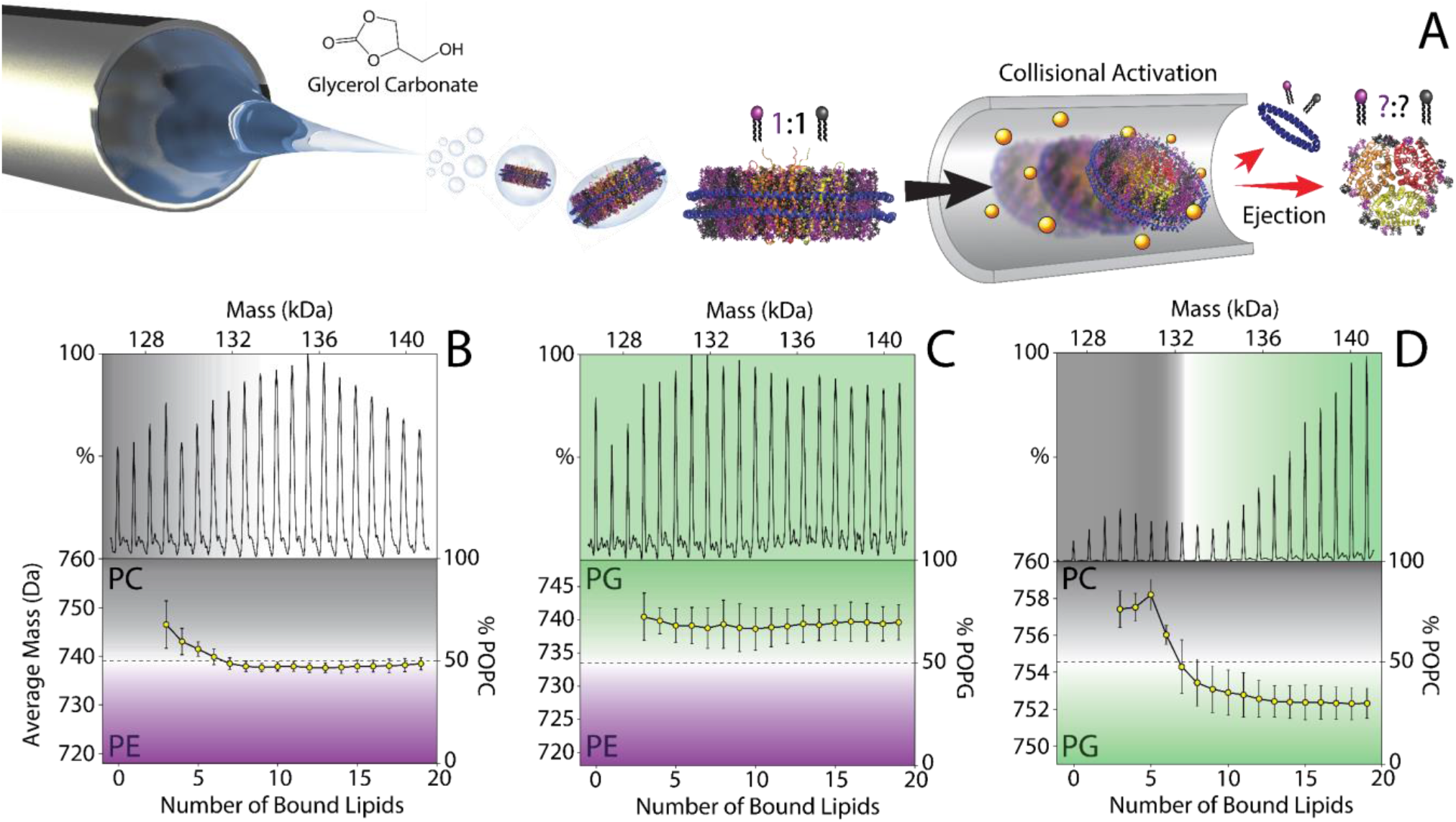
(A) Schematic for ejection of AmtB (PDB ID 4NH2)-lipid complexes from 50% PC:PE nanodiscs during native MS. Summed deconvolved mass spectra (*top*) of AmtB-lipid complexes ejected from (B) 50% PC:PE nanodiscs, (C) 50% PG:PE nanodiscs, and (D) 50% PC:PG nanodiscs. The corresponding average masses of bound lipids are indicated below. Black, green, and purple regions represent enrichment in PC, PG, and PE, respectively. White regions represent no lipid enrichment. The initial expected average lipid masses corresponding to 50% lipid mixtures are indicated by dashed lines. Average masses heavier or lighter than the dashed line show lipid enrichment. Figure 1D adapted from ref. 18. Copyright 2020 American Chemical Society.

We first performed native MS of AmtB-lipid complexes ejected from 50% PC:PE nanodiscs. Most bound lipids showed no enrichment, with average lipid masses almost exactly as expected for a 50:50 mixture of PC:PE. However, despite their similar chemical structures, the five lipids that were most tightly retained during ejection of AmtB had higher than expected average lipid masses, indicating slight enrichment in PC (Figure 1B). Thus, AmtB was not generally selective for PC over PE, except for a few bound lipids.

To further examine the lipid selectivity of AmtB, we ejected it from 50% PG:PE nanodiscs. Here, lipids bound to AmtB revealed around a 2:1 enrichment in PG over PE, which was consistent across all 19 lipid-bound states studied (Figure 1C). For example, the composition for 10 bound lipids was 66/34 ± 10% PG/PE. Thus, AmtB remodels mixed PG:PE bilayers to selectively bind PG lipids in a roughly 2:1 ratio.

After observing consistent PG enrichment and slight PC enrichment for a few selectively bound lipids, we compared the lipid enrichment with previous data from AmtB-lipid complexes ejected from 50% PC:PG nanodiscs (Figure 1D). Here, AmtB had distinct enrichment for two subsets of lipid-bound states. The first six lipids that were most tightly retained showed enrichment of PC. Conversely, the next 13 bound lipids showed enrichment of PG. Specifically, the bound lipid composition for AmtB bound to 3 and 19 lipids was 77/23 ± 9% PC/PG and 30/70 ± 8% PC/PG, respectively.

Collectively, these datasets reveal that AmtB selectively remodels its local lipid environment in two different ways. First, the most tightly retained lipids were enriched in PC, showing higher enrichment against PG but still detectable enrichment over PE. PE lipids only differ from PC lipids by three methyl groups, so these surprising results suggest that PC selectivity may be due to a few very specific binding interactions. Interestingly, PC binding has been shown to enhance the binding of AmtB to the regulatory protein, GlnK,^29^ so these PC binding sites may affect AmtB interactions. Second, AmtB was enriched in PG in both PG:PE and PC:PG mixtures, agreeing with previously established functional and stabilizing roles of PG for AmtB.^8, 19^ No selective enrichment of PE was observed for either lipid mixture, so PE seems to play the role of a generic neutral lipid that participates non-selectively in the annular belt.

### Ejection of AmtB from Three-Component Lipid Nanodiscs Confirms PC Selectivity

To probe lipid enrichment in a more complex lipid environment, we ejected AmtB from nanodiscs with a ternary mixture of all three lipids (Figure S-3 and S-4). Because there are three different mass components, we unfortunately cannot calculate an exact lipid composition from the average lipid mass. However, we can make predictions based on the binary data above and rule out combinations that are not consistent with the measured average masses.

The average masses of most bound lipids were consistent with somewhere between no enrichment (a 1:1:1 mixture) and a 2:1 PG enrichment (2:1:1 PG:PC:PE) (Figure 2). Thus, we can rule out PE or PC enrichment, which would significantly alter the average lipid masses. These data are consistent with overall enrichment of PG in ternary nanodiscs, but we cannot confidently measure the degree of enrichment. Interestingly, the few most tightly retained lipids showed a gradual shift to heavier masses, supporting a likely enrichment of PC observed in binary data. Overall, the data from AmtB in ternary nanodiscs is consistent with AmtB being selective for PC for a few tightly retained lipids and selective for PG overall. However, the PC enrichment appears less substantial than in PC:PG nanodiscs, suggesting that the presence of PE dampens the lipid selectivity of AmtB in ternary mixtures.

**Figure 2.**
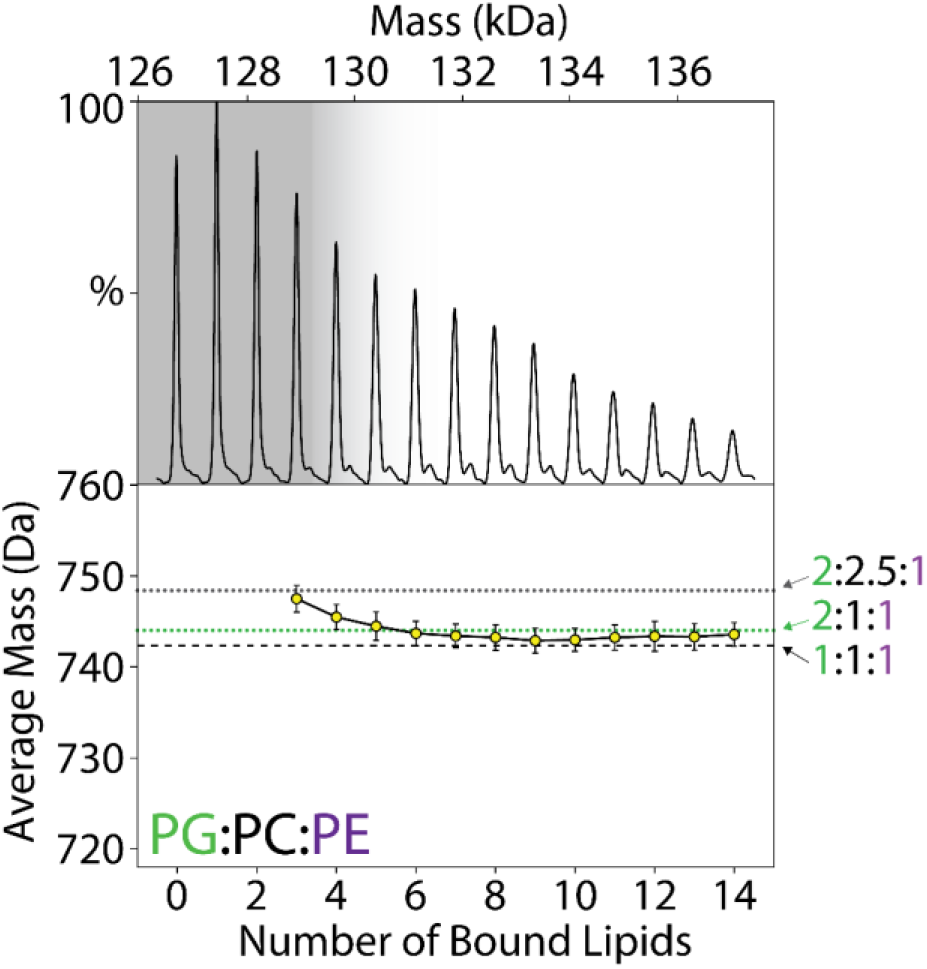
Summed deconvolved mass spectrum of AmtB-lipid complexes ejected from 1:1:1 PG:PC:PE nanodiscs. The corresponding average masses of bound lipids are indicated below. Dashed and dotted lines are annotated with possible lipid compositions corresponding to no enrichment (1:1:1), PG enrichment (2:1:1), and PC enrichment (2:2.5:1). Grey shading indicates possible PC enrichment.

### Detergent Extraction for Orthogonal Characterization of Lipid Selectivity

Ejection of AmtB-lipid complexes from heterogeneous nanodiscs allows us to probe selective lipid remodeling for a wide range of lipid-bound states,^18^ but it can be hard to relate the most tightly retained lipids upon gas-phase ejection with the most tightly bound lipids in solution. Robinson and coworkers demonstrated that lipid binding can be sensitive to electrospray polarity.^30^ However, they observed that AmtB showed little change in lipid binding between electrospray polarities. Recent studies from Prell and coworkers^28^ propose that different lipid head groups can dissociate differently at high levels of collisional activation due to differences in gas-phase basicity. Although we did not observe lipid compositions expected from this type of gas-phase dissociation, different gas-phase binding strengths could bias which lipids are retained.

To compare our gas-phase results with solution-phase binding, we developed a new approach to rapidly extract AmtB-lipid complexes in solution rather than the gas phase (Figure 3A). Previous studies show that the exchange between bound lipids and detergent molecules can be used to distinguish non-annular lipid interactions.^31–32^ Thus, we added detergents to break up the nanodisc and displace non- or weakly-interacting lipids around AmtB. We tested tetraethylene glycol monooctyl ether (C8E4), Triton X-100, and lauryl dimethyl amine oxide (LDAO) because they require low collisional activation to dissociate from membrane proteins.^33–34^ Saccharide detergents are common for membrane protein solubilization, so we also tested octyl glucoside (OG).

**Figure 3.**
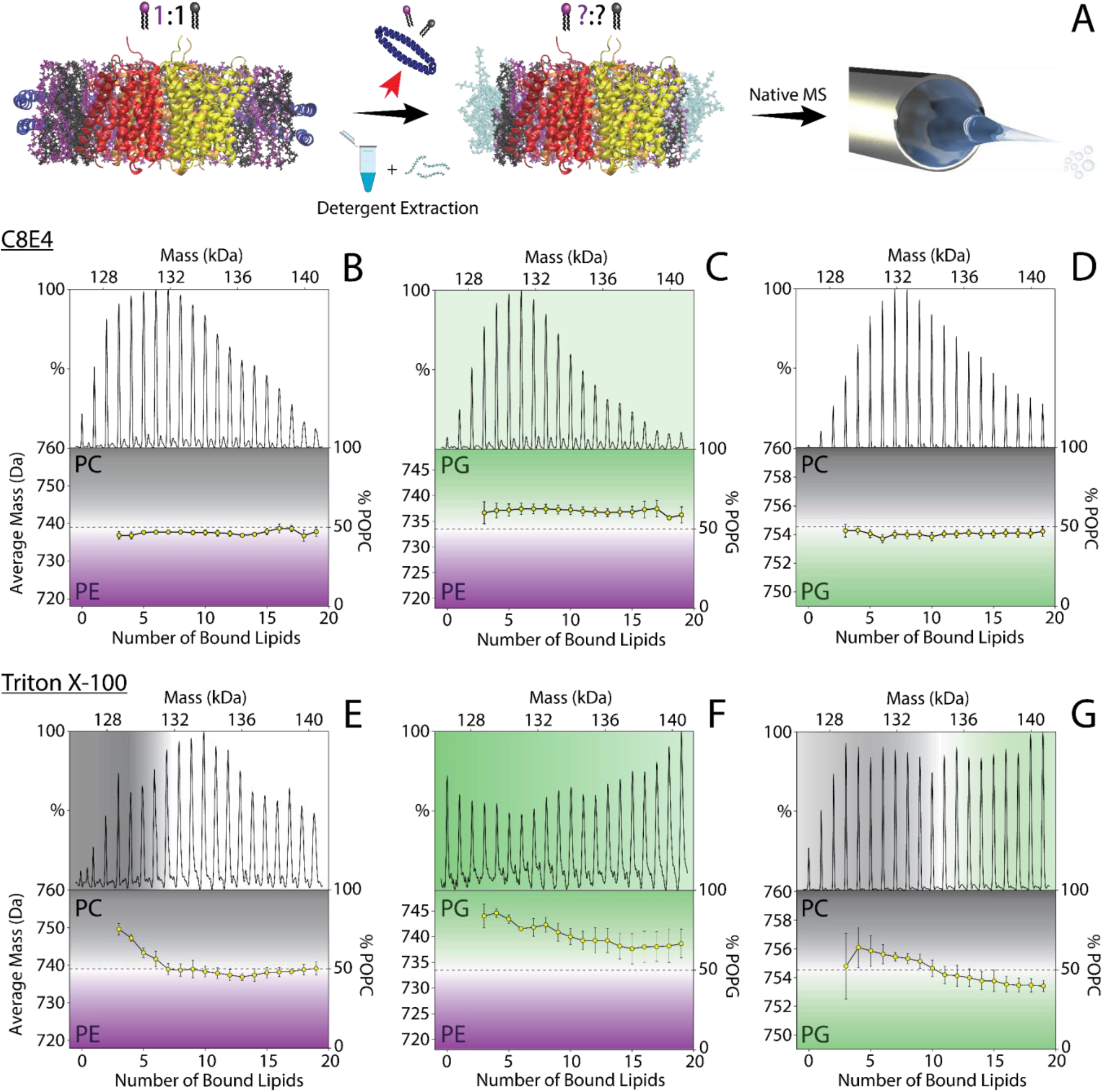
(A) Schematic for detergent extraction of AmtB-lipid complexes from binary lipid nanodiscs for native MS. Summed deconvolved mass spectra of AmtB-lipid complexes extracted with C8E4 (B–D) or Triton X-100 (E–G) from 50% (B, E) PC:PE nanodiscs, (C, F) PG:PE nanodiscs, and (D, G) PC:PG nanodiscs. The corresponding average masses of bound lipids are plotted below the deconvolved mass spectra. Black, green, and purple regions represent enrichment in PC, PG, and PE, respectively. White regions represent no lipid enrichment. The initial expected average lipid masses corresponding to 50% lipid mixtures are indicated by dashed lines.

To test their capacity for detergent extraction of AmtB from nanodiscs, different incubation times and detergent concentrations were screened. With C8E4 and Triton X-100, AmtB could easily be captured with both high and low numbers of bound lipids following minimal incubation times. Conversely, AmtB could only be captured with high numbers of bound lipids following incubation with LDAO and OG. Higher concentrations and longer incubation times helped with isolating lower numbers of bound lipids, but we continued with only C8E4 and Triton X-100. The final concentration of detergent was limited to 2× the critical micelle concentration (CMC) to improve mass spectra quality. Native MS revealed that AmtB is extracted into mixed lipid-detergent complexes devoid of MSP (Figure S-5–S-8).

With C8E4, AmtB extracted from 50% PC:PE nanodiscs had almost no enrichment (Figure 3B). AmtB from 50% PG:PE was slightly enriched in PG, statistically similar (p<0.05 by T-test) to gas-phase ejection studies (Figure 3C). Finally, AmtB extracted from 50% PC:PG also showed little to no enrichment (Figure 3D). Overall, although the preference for PG over PE was retained, the PC selectivity of AmtB observed with gas-phase ejection was mostly absent when extracted into C8E4 detergent micelles.

We then probed lipid enrichment when extracting with Triton X-100. Here, AmtB extracted from 50% PC:PE nanodiscs had PC enrichment for the five most-tightly bound lipids and no enrichment for the next 14 lipids (Figure 3E), similar to results from gas-phase ejection. For 50% PG:PE nanodiscs, all lipids were enriched in PG with lower numbers of lipids showing progressively more enrichment (Figure 3F). Finally, AmtB extracted from 50% PC:PG nanodiscs had distinct enrichment for two subsets of lipids (Figure 3G). The first subset of tightly bound lipids was enriched in PC, and the less tightly bound subset was enriched in PG. In all three cases, Triton X-100 extraction qualitatively matched the lipid selectivity observed with gas-phase ejection.

Finally, we extracted AmtB from 1:1:1 PG:PC:PE nanodiscs (Figure S-9 and S-10). With C8E4, all lipids showed a similar average lipid mass, consistent with a constant lipid enrichment in PG but a loss of selective PC binding for tightly bound lipids (Figure 4A). For Triton X-100, lipids showed a gradually heavier average mass for more tightly bound lipids, indicating likely PC enrichment for lower numbers of bound lipids (Figure 4B). Thus, the lipid selectivity for AmtB extracted from ternary nanodiscs with Triton X-100 qualitatively agrees with both the Triton X-100 dataset from binary nanodiscs and with gas-phase ejection from nanodiscs. Together, these results indicate a broad selectivity for PG lipids, a few tight binding sites for PC, and limited preference for PE.

**Figure 4.**
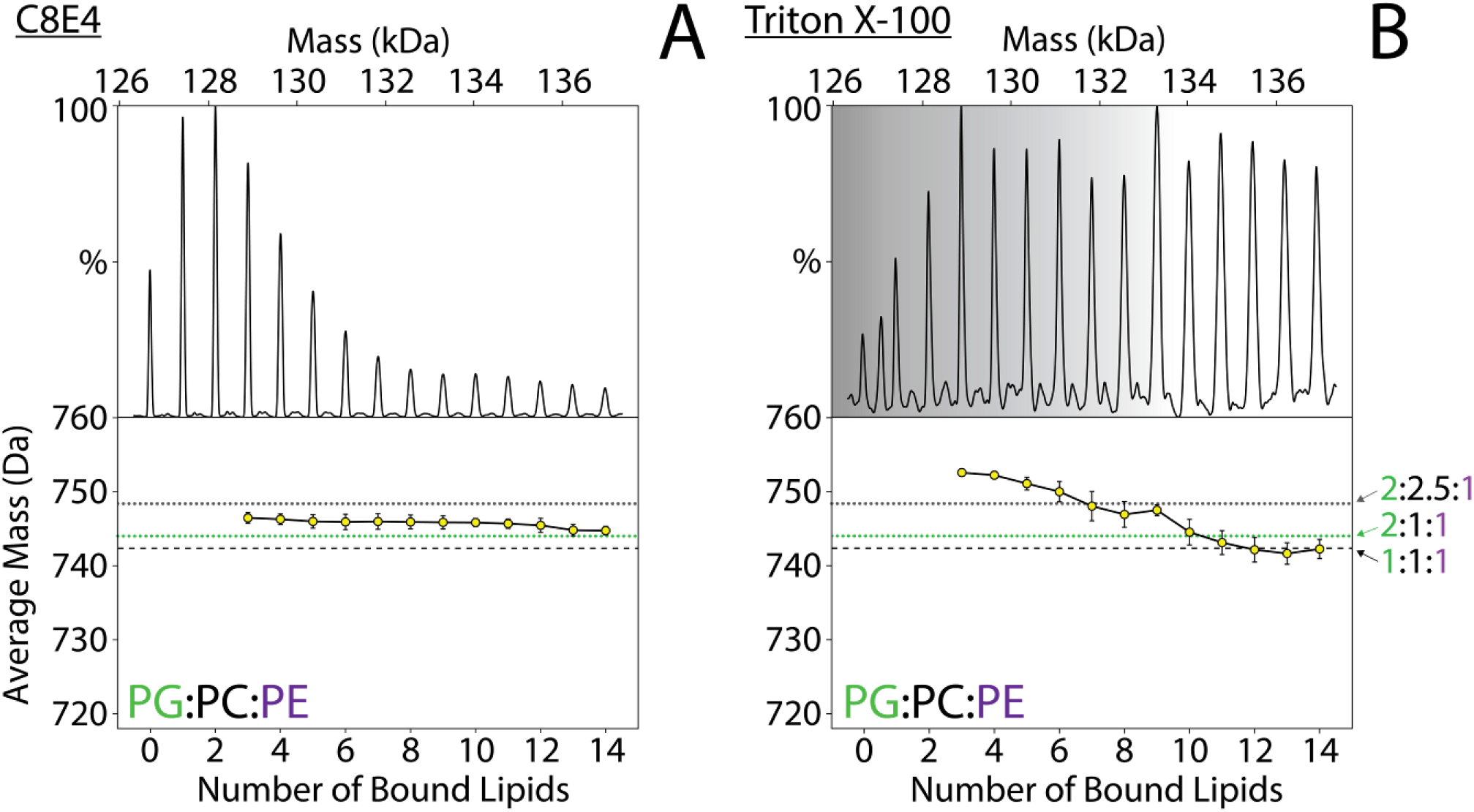
Summed deconvolved mass spectra of AmtB-lipid complexes extracted from 1:1:1 PG:PC:PE lipid nanodiscs using (A) C8E4 and (B) Triton X-100. The corresponding average masses of bound lipids are indicated below. Dashed and dotted lines are annotated with possible lipid compositions corresponding to no enrichment, PG enrichment, and PC enrichment. Grey shading indicates possible PC enrichment.

### Lipid Selectivity of AqpZ in Nanodiscs

To investigate a second membrane protein system, we assembled AqpZ into nanodiscs with 50% PC:PE, PG:PE, and PC:PG and performed native MS on AqpZ that was either ejected in the gas phase or through solution-phase extraction (Figure S-11–S-13). Unfortunately, extraction with Triton X-100 yielded spectra that were poorly resolved, so we substituted LDAO as a second detergent to compare with C8E4.

Comparing the average lipid mass values from gas-phase ejection, LDAO extraction, and C8E4 extraction (Figure 5), there were strong qualitative similarities between the gasphase ejection and LDAO extraction. Both showed preferences for PC over PE (Figure 5A, D), PG over PE (Figure 5B, E), and PC over PG (Figure 5C, F). The average enrichment for PG/PE was quantitatively different between the LDAO and gas-phase ejection, but the differences were statistically insignificant for the other lipid combinations.

**Figure 5.**
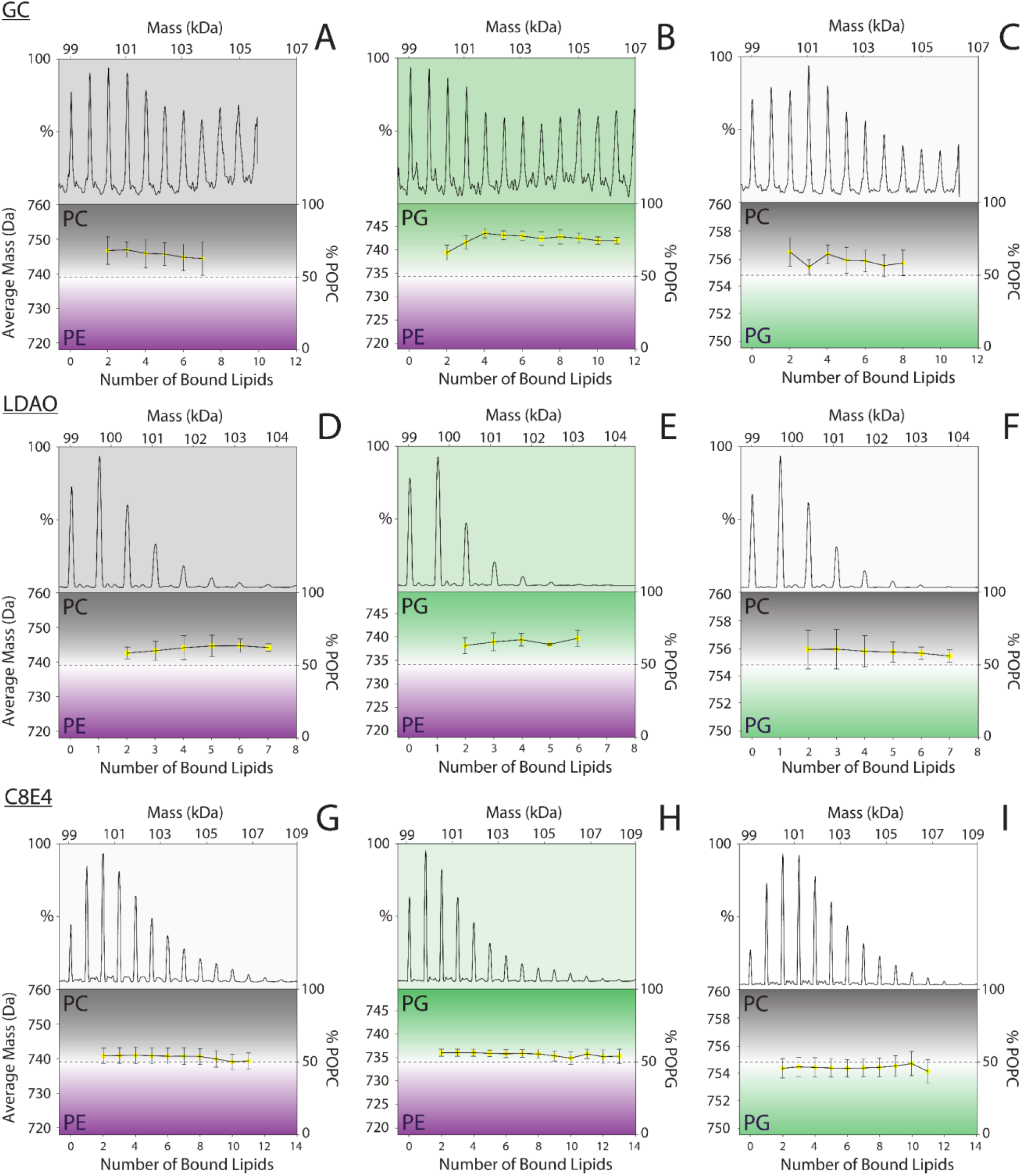
Summed deconvolved mass spectra of AqpZ-lipid complexes ejected (A–C), extracted with LDAO (D–F), or extracted with C8E4 (G–I) from (A, D, G) 50% PC:PE nanodiscs, (B, E, H) 50% PG:PE nanodiscs, or (C, F, I) 50% PC:PG nanodiscs. The corresponding average masses of bound lipids are plotted below the deconvolved mass spectra. Black, green, and purple regions represent enrichment in PC, PG, and PE, respectively. White regions represent no lipid enrichment. The expected average lipid masses corresponding to 50% lipid mixtures are indicated by dashed lines. Average masses heavier or lighter than the dashed line show lipid enrichment.

In contrast, AqpZ extracted with C8E4 had no selectivity for PC over PE (Figure 5G), a slight preference for PG over PE (Figure 5H), and no preference for PG over PC (Figure 5I). Statistically significant differences were observed between C8E4 and gas-phase ejection for all lipid combinations. Interestingly, all detergent and ejection conditions showed more lipids retained on AqpZ when PC was included in the mixture. The lowest total numbers of bound lipids were observed with PG:PE.

Comparing the selectivity profiles of AqpZ with AmtB for different approaches (Figures 1, 3, and 5), there was consistent qualitative agreement between gas-phase ejection and either Triton X-100 or LDAO across all three lipid mixtures. With C8E4, there was consistent agreement in selectivity for mixtures of PG:PE. However, C8E4 extraction did not completely preserve the PC selectivity observed with the other detergents and gas-phase ejection.

The differences observed between C8E4, LDAO, and Triton X-100 reinforce that there is a delicate balance for detergents between compatibility with native MS, solubilization effectiveness, and disruption of lipid interactions. These factors should be taken into careful consideration when selecting detergents for extracting membrane proteins. An interesting approach would be to use novel oligoglycerol detergents^35^ to fine-tune the propensity for delipidation to study different types of protein-lipid interactions.

Looking broadly at the data, AqpZ and AmtB shared some similar lipid preferences but also showed some notable differences. Both membrane proteins had a preference for PG over PE lipids that was relatively constant across the number of bound lipids measured here. The consistent preference for anionic PG over zwitterionic PE likely reveals several charge-specific lipid binding sites with modest but not absolute selectivity. However, interactions with PC were different between the two proteins. AmtB had little selectivity for PC except for a small number of very specific binding sites. AqpZ showed a more consistent selectivity for PC over both PE and PG, lacking the bimodal binding present with AmtB. Overall, these results provide an initial picture of the relative selectivity of two membrane proteins for bilayers with controlled mixtures of lipids.

## CONCLUSIONS

Mounting evidence is revealing the vital role of lipids in membrane protein structure and function.^13, 36–39^ High-resolution biophysical techniques can provide detailed structures of membrane proteins in stable complexes with nonannular lipids. However, many lipid interactions are transient and heterogeneous, and it is challenging to measure how membrane proteins selectively remodel their surrounding lipid environment to bind these annular lipids. Here, we demonstrated that native MS can be used to probe lipid remodeling by membrane proteins in heterogeneous lipid nanodiscs and distinguish lipid selectivity for a larger number of bound lipids.

Gas-phase ejection and solution-phase detergent extraction for native MS provide orthogonal approaches for investigating how AmtB and AqpZ remodel their local lipid environment in two- and three-component nanodiscs. Both approaches reveal that AmtB is broadly enriched in bound PG, has a few tightly bound PC lipids, but shows no selective enrichment in PE. In contrast, AqpZ prefers either PC or PG over PE and PC over PG.

Detergent extraction provides a quick, simple, and complementary approach for investigating membrane protein-lipid interactions but requires careful selection of detergents. Triton X-100 and LDAO extraction preserved enrichment of PC and PG lipids, but C8E4 extraction showed little to no enrichment, demonstrating that detergents can selectively alter membrane protein-lipid interactions when extracting from lipid bilayers. Overall, these two approaches are broadly applicable for distinguishing and quantifying membrane protein-lipid selectivity, providing detailed insights into the biophysics of membrane protein-lipid interactions in heterogeneous lipid bilayers.

## ASSOCIATED CONTENT

### Supporting Information

Native mass spectra for AmtB and AqpZ nanodiscs. Analysis for simulated mass spectra of AmtB-lipid complexes. This material is available free of charge via the Internet at http://pubs.acs.org.

## AUTHOR INFORMATION

### Notes

The authors declare no competing financial interest.

## ACKNOWLEDGMENTS

The authors thank Maria Reinhardt-Szyba, Kyle Fort, and Alexander Makarov at Thermo Fisher Scientific for support on the Q-Exactive HF UHMR instrument. The pMSP1E3D1 plasmid was a gift from Stephen Sligar (Addgene plasmid no. 20066). This work was funded by the National Institute of General Medical Sciences and National Institutes of Health (Grant R35 GM128624). The content is solely the responsibility of the authors and does not necessarily represent the official views of the NIH.

## For Table of Contents Only

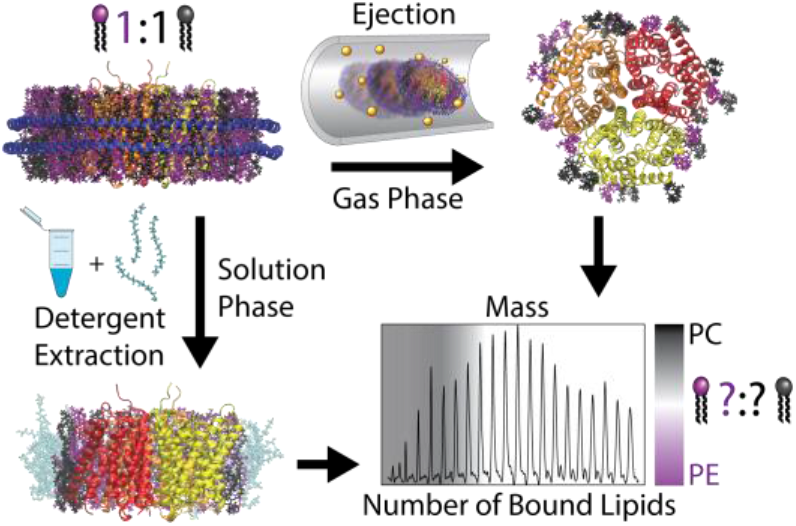

## Notes

### Competing Interest Statement

The authors have declared no competing interest.

### Summary of Updates

This manuscript has been updated in response to two rounds of reviewer comments.

