## Supporting Information for "Investigating the Lipid Selectivity of Membrane Proteins in Heterogeneous Nanodiscs"

### SUPPLEMENTAL FIGURES

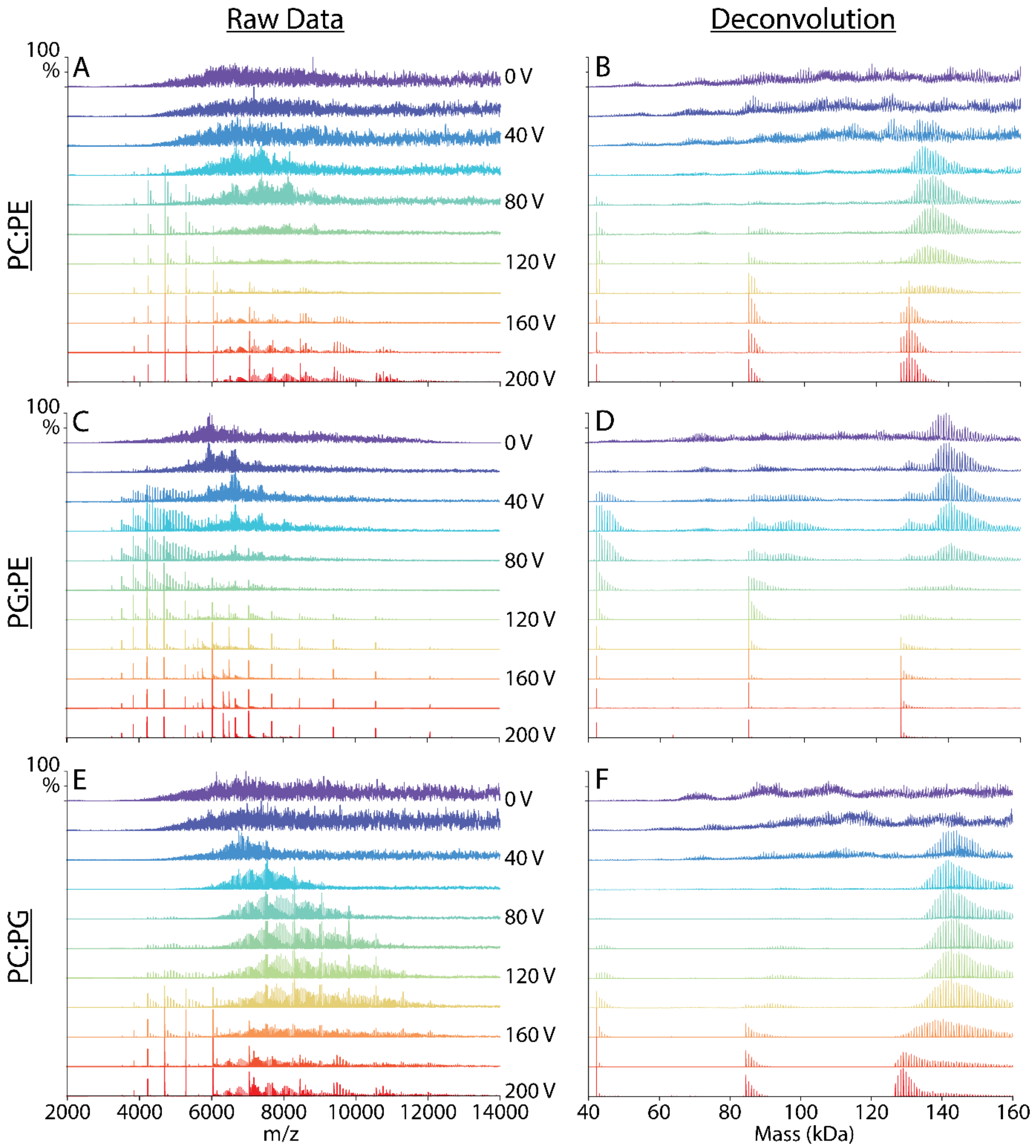

**Figure S-1.** Representative mass spectra (A, C, E) and zero-charge deconvolved spectra (B, D, F) of AmtB-lipid complexes ejected from 50% PC:PE (A, B), 50% PG:PE (C, D), and 50% PC:PG (E, F) binary lipid nanodiscs. Spectra are shown for increasing collision voltage from 0 V (purple) to 200 V (red) in 20 V increments.

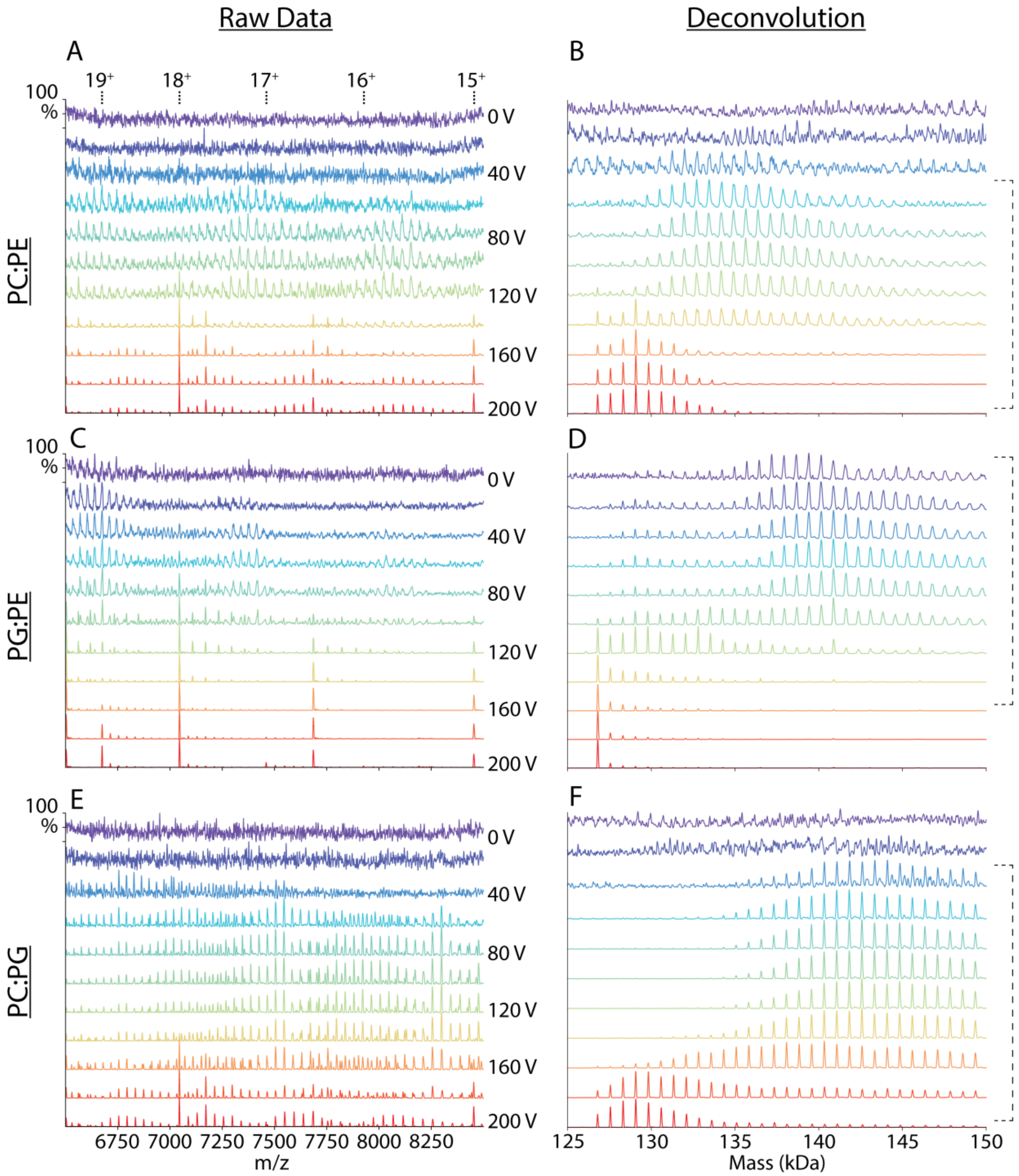

**Figure S-2.** Zoomed regions for representative mass spectra (A, C, E) and zero-charge deconvolved spectra (B, D, F) from Figure S-1. Charge states of AmtB are annotated for the raw data. Dashed brackets indicate voltages used for data analysis.

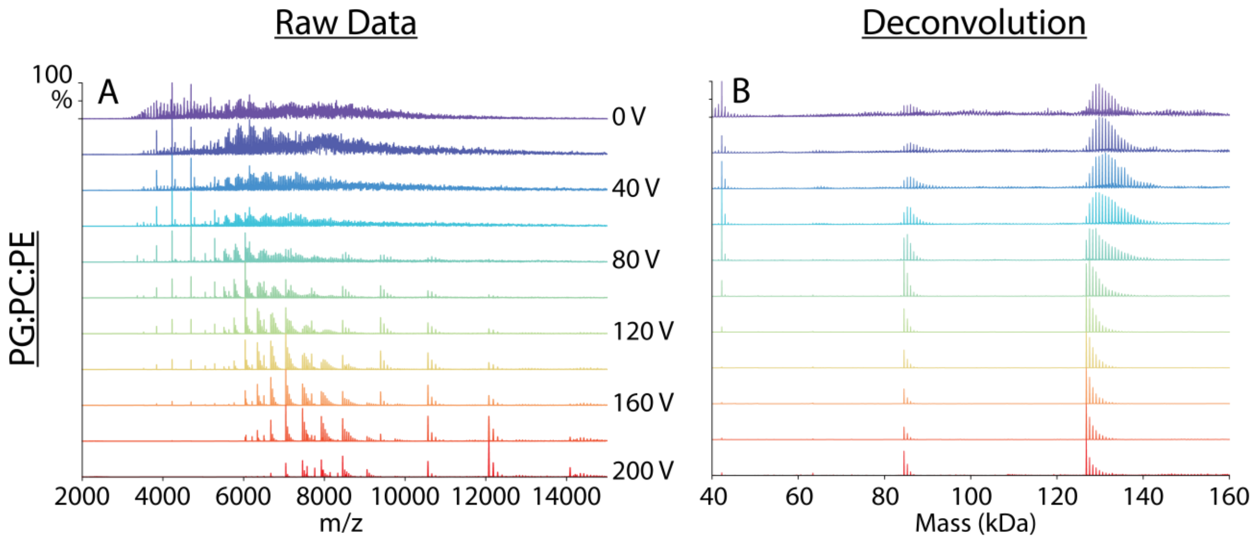

**Figure S-3.** Representative mass spectrum (A) and zero-charge deconvolved spectrum (B) of AmtB-lipid complexes ejected from 1:1:1 PG:PC:PE ternary lipid nanodiscs. Spectra are shown for increasing collision voltage from 0 V (purple) to 200 V (red) in 20 V increments.

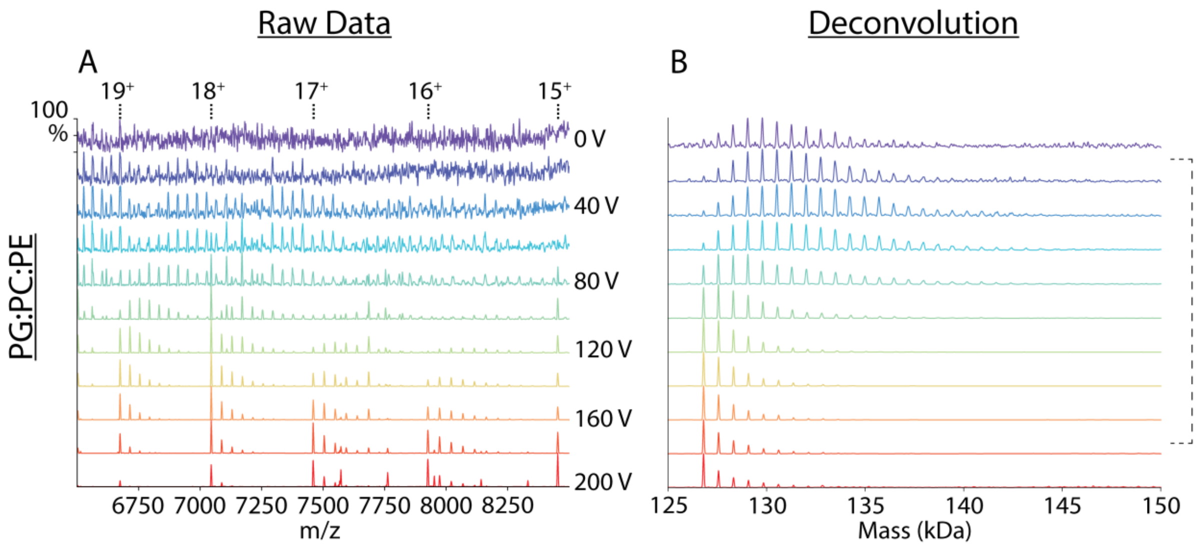

**Figure S-4.** Zoomed regions for representative mass spectrum (A) and zero-charge deconvolved spectrum (B) from Figure S-3. Charge states of AmtB are annotated for the raw data. Dashed brackets indicate voltages used for data analysis.

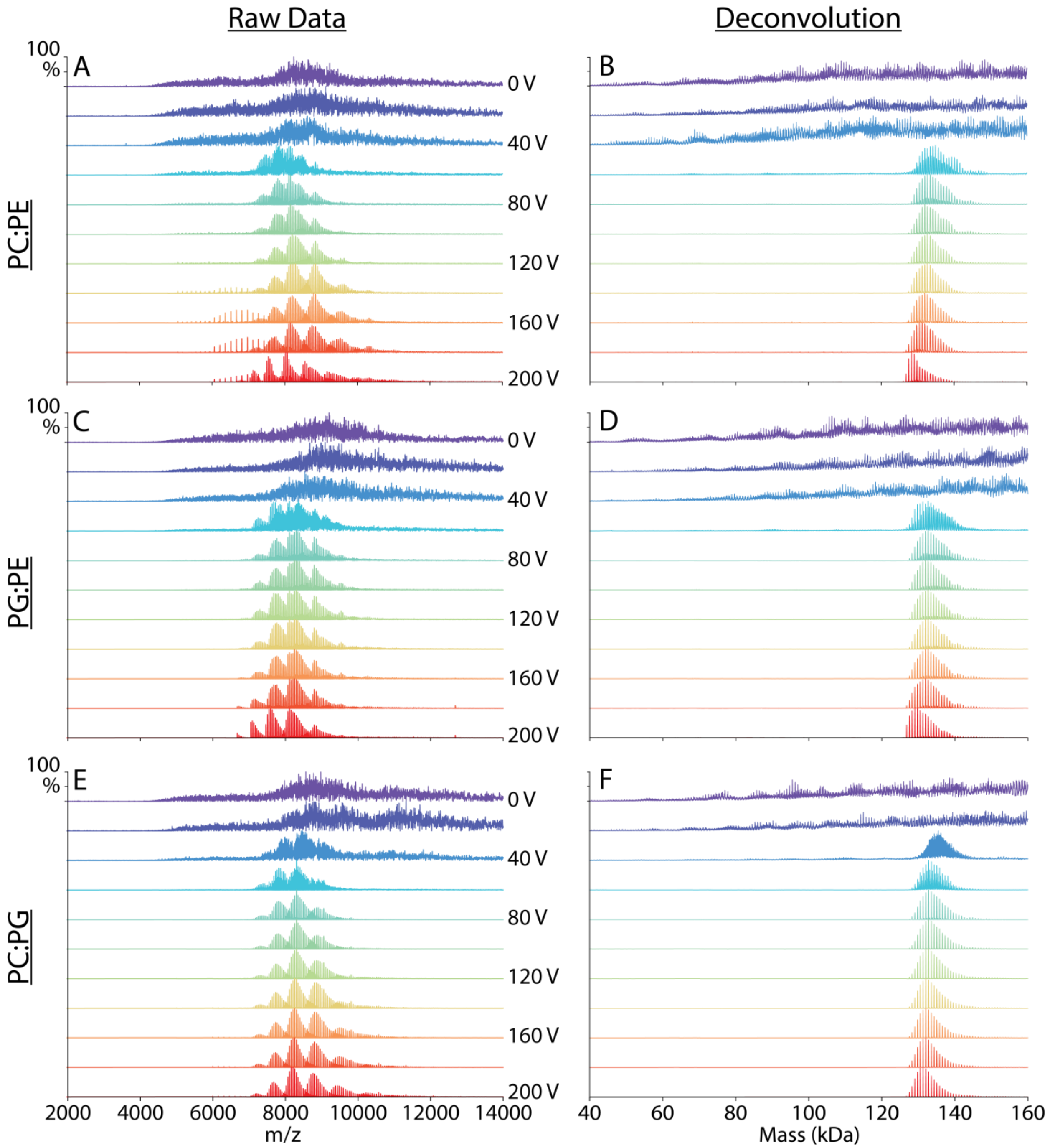

**Figure S-5.** Representative mass spectra (A, C, E) and zero-charge deconvolved spectra (B, D, F) of AmtB-lipid complexes extracted with C8E4 detergent from 50% PC:PE (A, B), 50% PG:PE (C, D), and 50% PC:PG (E, F) binary lipid nanodiscs. Spectra are shown for increasing collision voltage from 0 V (purple) to 200 V (red) in 20 V increments.

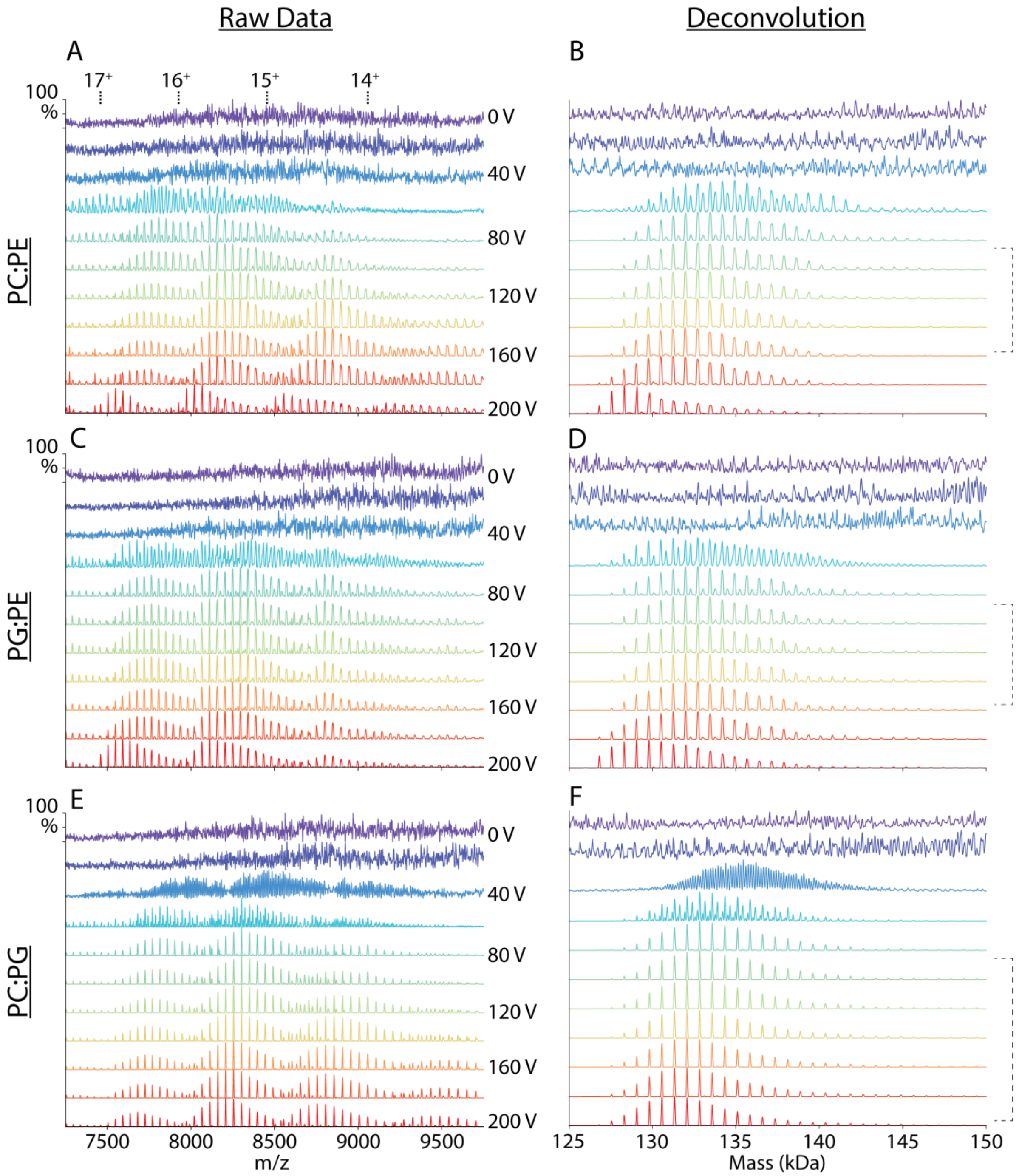

**Figure S-6.** Zoomed regions for representative mass spectra (A, C, E) and zero-charge deconvolved spectra (B, D, F) from Figure S-5. Charge states of AmtB are annotated for the raw data. Dashed brackets indicate voltages used for data analysis.

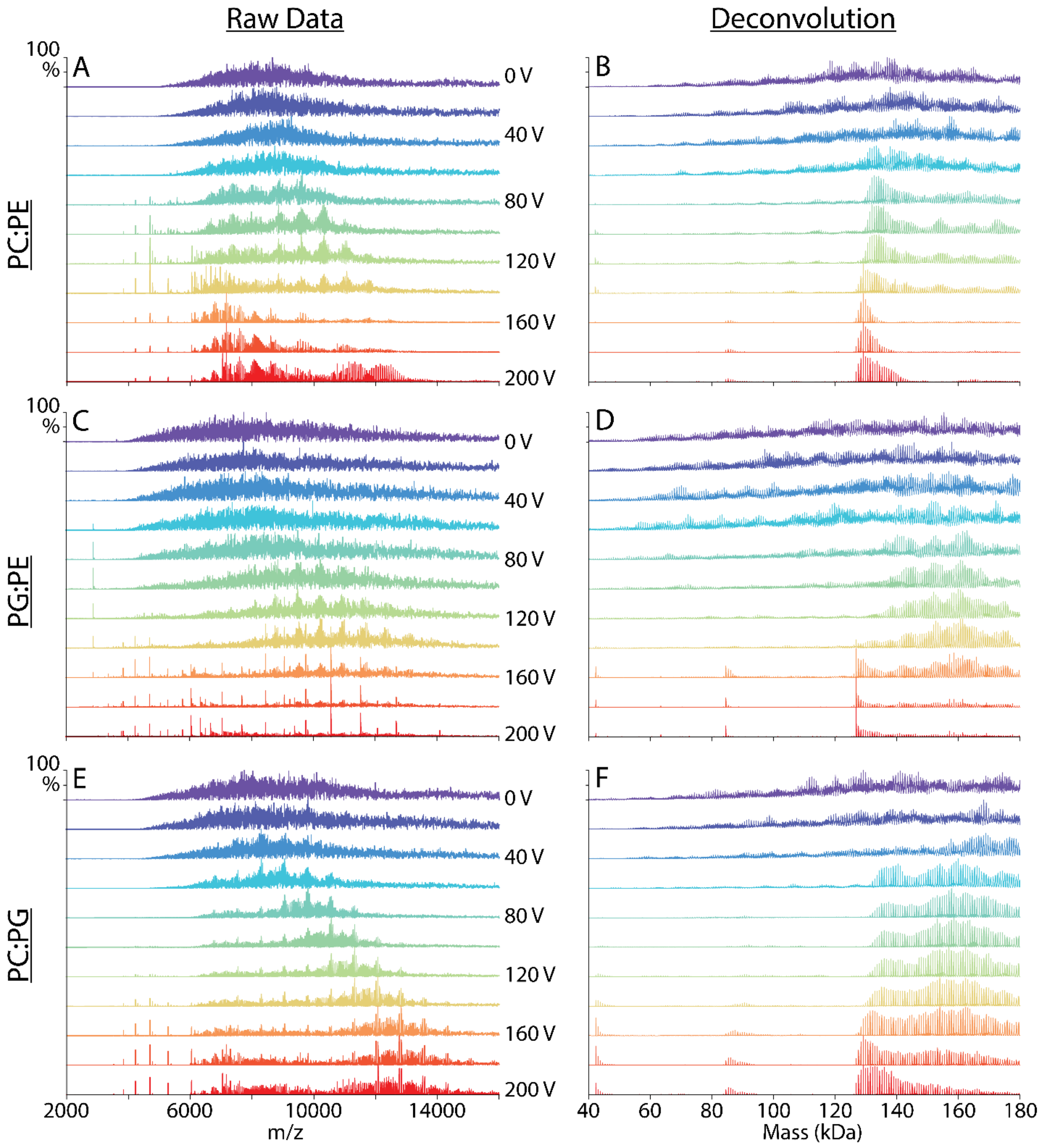

**Figure S-7.** Representative mass spectra (A, C, E) and zero-charge deconvolved spectra (B, D, F) of AmtB-lipid complexes extracted with Triton X-100 detergent from 50% PC:PE (A, B), 50% PG:PE (C, D), and 50% PC:PG (E, F) binary lipid nanodiscs. Spectra are shown for increasing collision voltage from 0 V (purple) to 200 V (red) in 20 V increments.

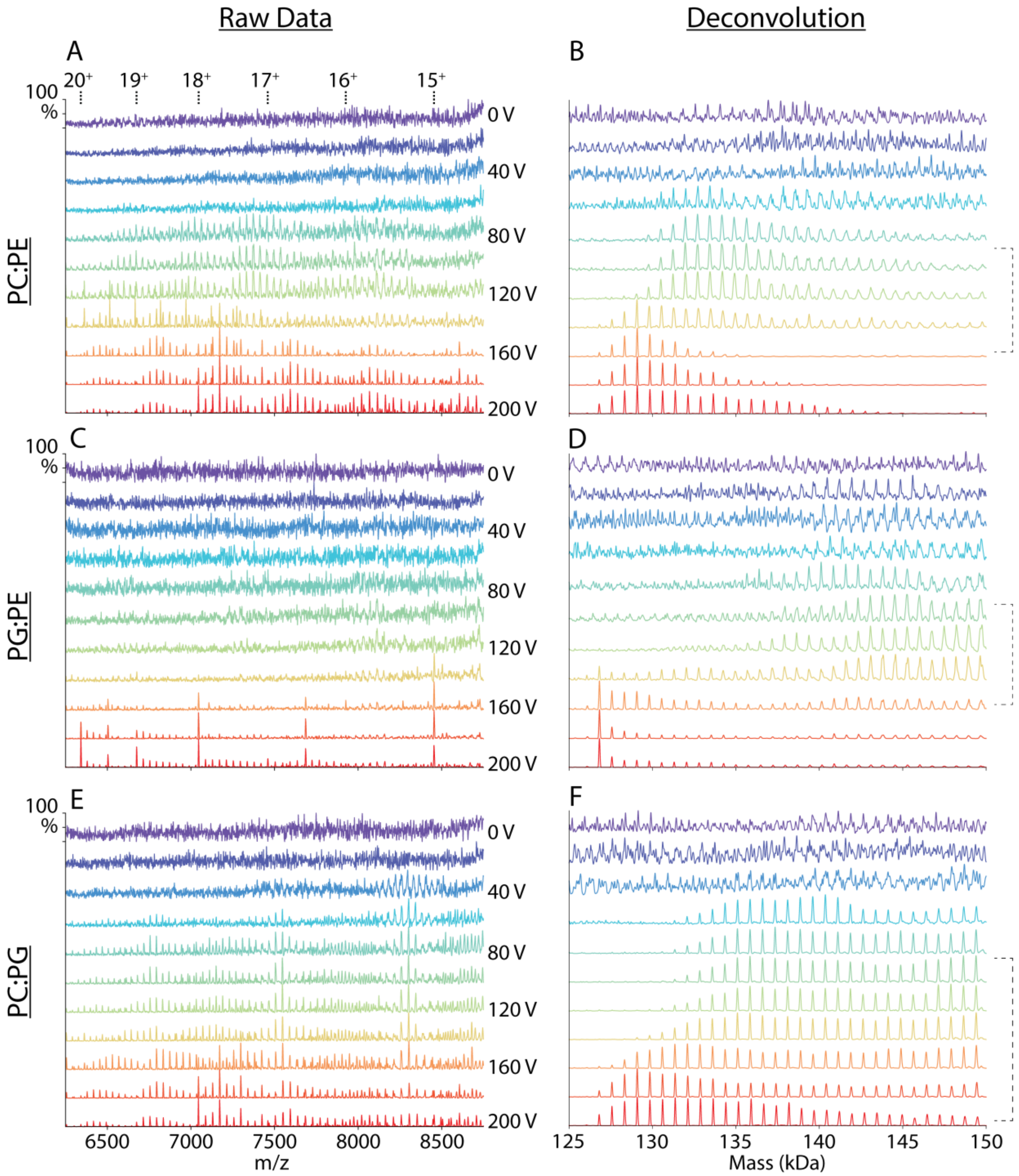

**Figure S-8.** Zoomed regions for representative mass spectra (A, C, E) and zero-charge deconvolved spectra (B, D, F) from Figure S-7. Charge states of AmtB are annotated for the raw data. Dashed brackets indicate voltages used for data analysis.

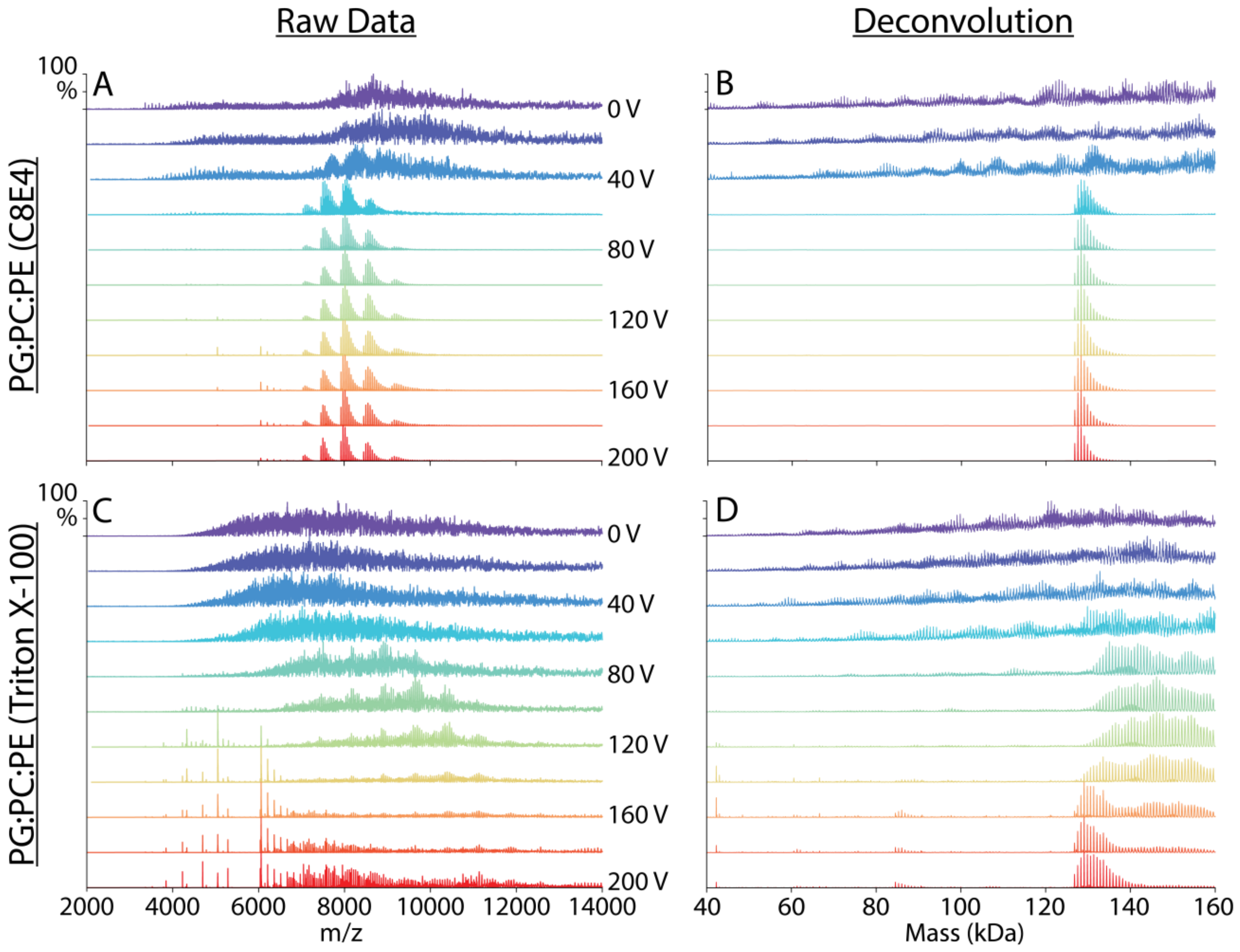

**Figure S-9.** Representative mass spectra (A, C) and zero-charge deconvolved spectra (B, D) of AmtB-lipid complexes extracted with C8E4 (A, B) and Triton X-100 (C, D) from 1:1:1 PG:PC:PE ternary lipid nanodiscs. Spectra are shown for increasing collision voltage from 0 V (purple) to 200 V (red) in 20 V increments.

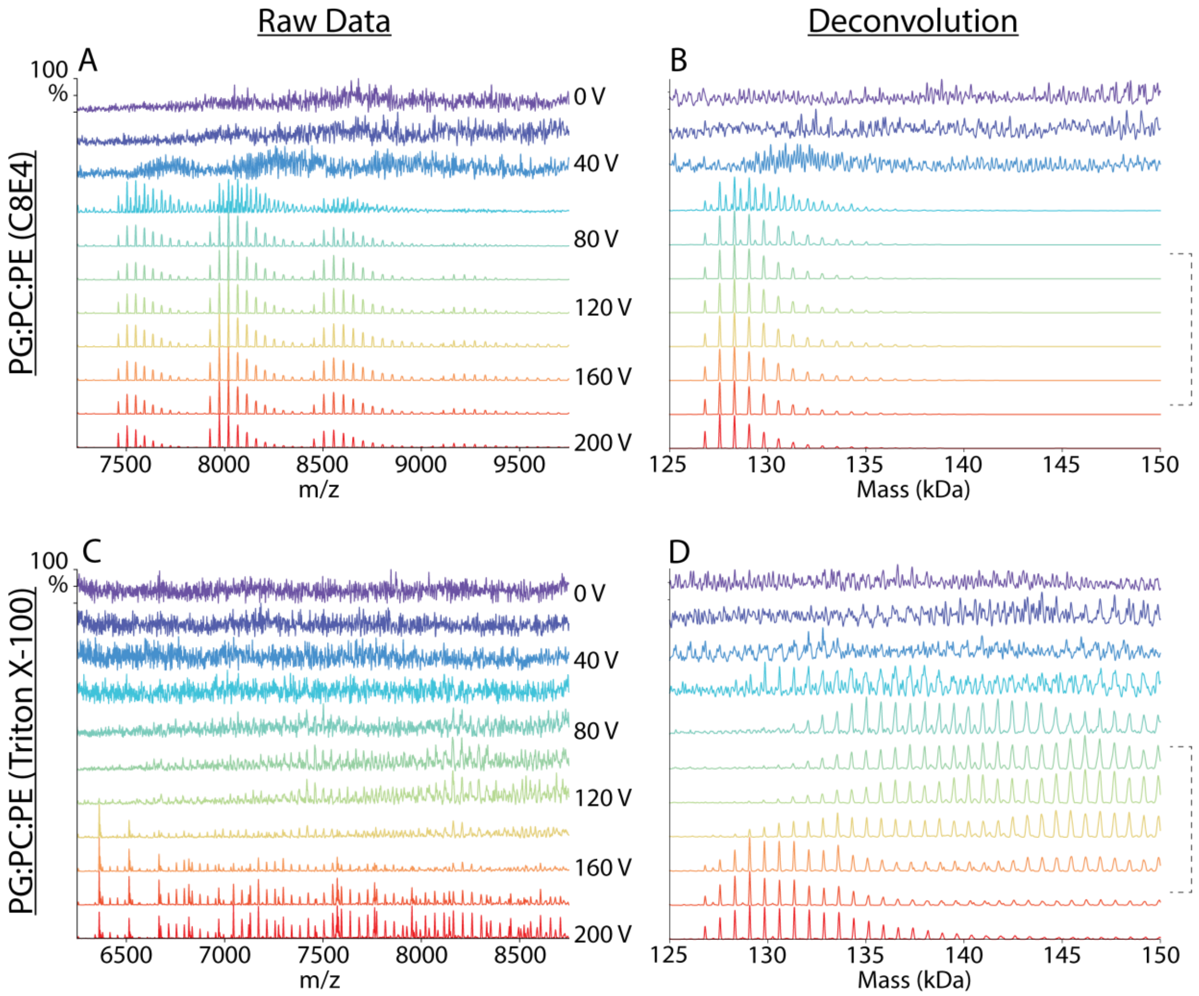

**Figure S-10.** Zoomed regions for representative mass spectra (A, C) and zero-charge deconvolved spectra (B, D) from Figure S-9. Dashed brackets indicate voltages used for data analysis.

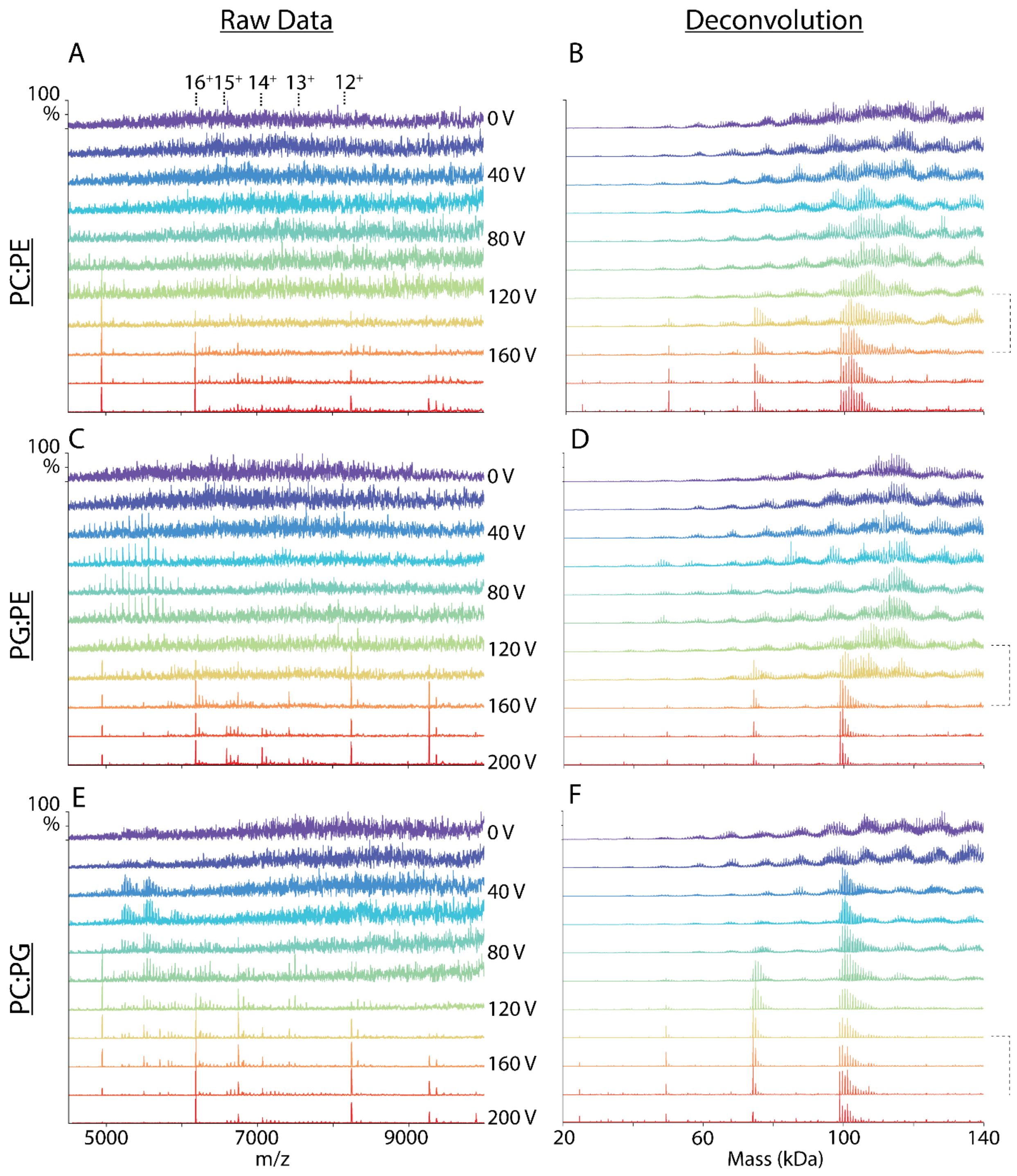

**Figure S-11.** Representative mass spectra (A, C, E) and zero-charge deconvolved spectra (B, D, F) of AqpZ-lipid complexes ejected from 50% PC:PE (A, B), 50% PG:PE (C, D), and 50% PC:PG (E, F) binary lipid nanodiscs. Spectra are shown for increasing collision voltage from 0 V (purple) to 200 V (red) in 20 V increments. Charge states of AqpZ are annotated for the raw data. Dashed brackets indicate voltages used for data analysis.

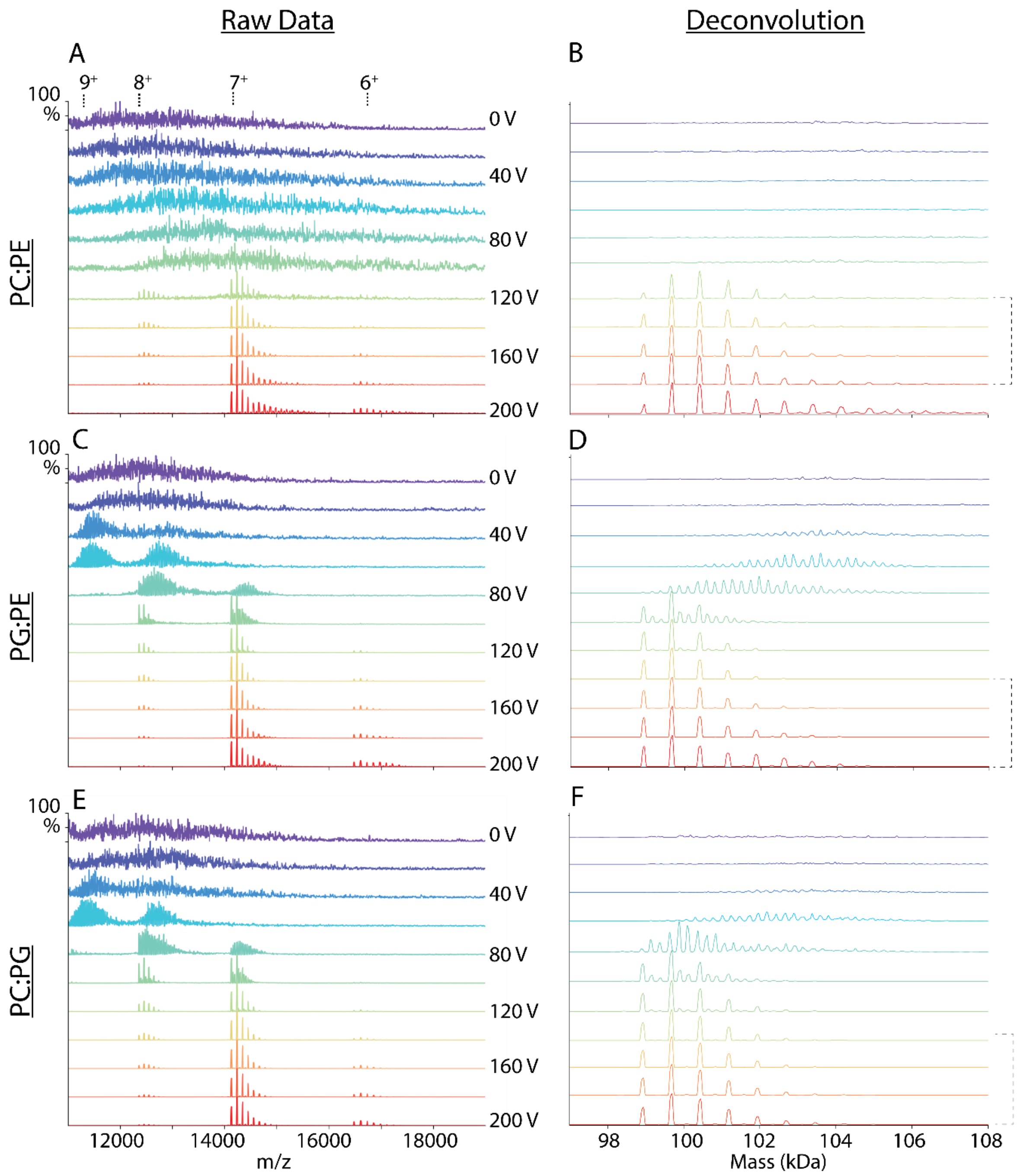

**Figure S-12.** Zoomed regions for representative mass spectra (A, C, E) and zero-charge deconvolved spectra (B, D, F) of AqpZ-lipid complexes extracted with LDAO detergent from 50% PC:PE (A, B), 50% PG:PE (C, D), and 50% PC:PG (E, F) binary lipid nanodiscs. Spectra are shown for increasing collision voltage from 0 V (purple) to 200 V (red) in 20 V increments. Charge states of AqpZ are annotated for the raw data. Dashed brackets indicate voltages used for data analysis.

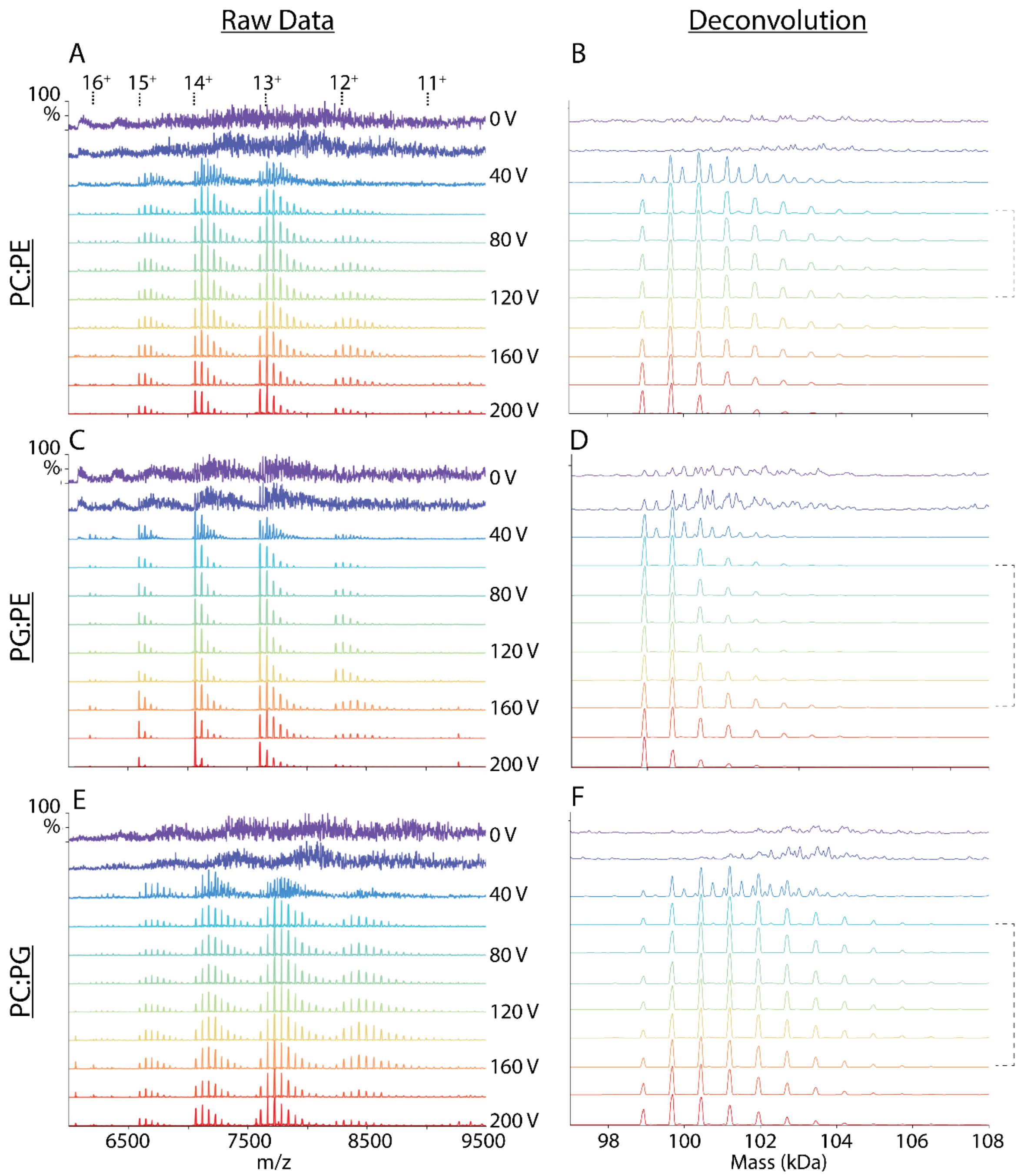

**Figure S-13.** Zoomed regions for representative mass spectra (A, C, E) and zero-charge deconvolved spectra (B, D, F) of AqpZ-lipid complexes extracted with C8E4 detergent from 50% PC:PE (A, B), 50% PG:PE (C, D), and 50% PC:PG (E, F) binary lipid nanodiscs. Spectra are shown for increasing collision voltage from 0 V (purple) to 200 V (red) in 20 V increments. Charge states of AqpZ are annotated for the raw data. Dashed brackets indicate voltages used for data analysis.

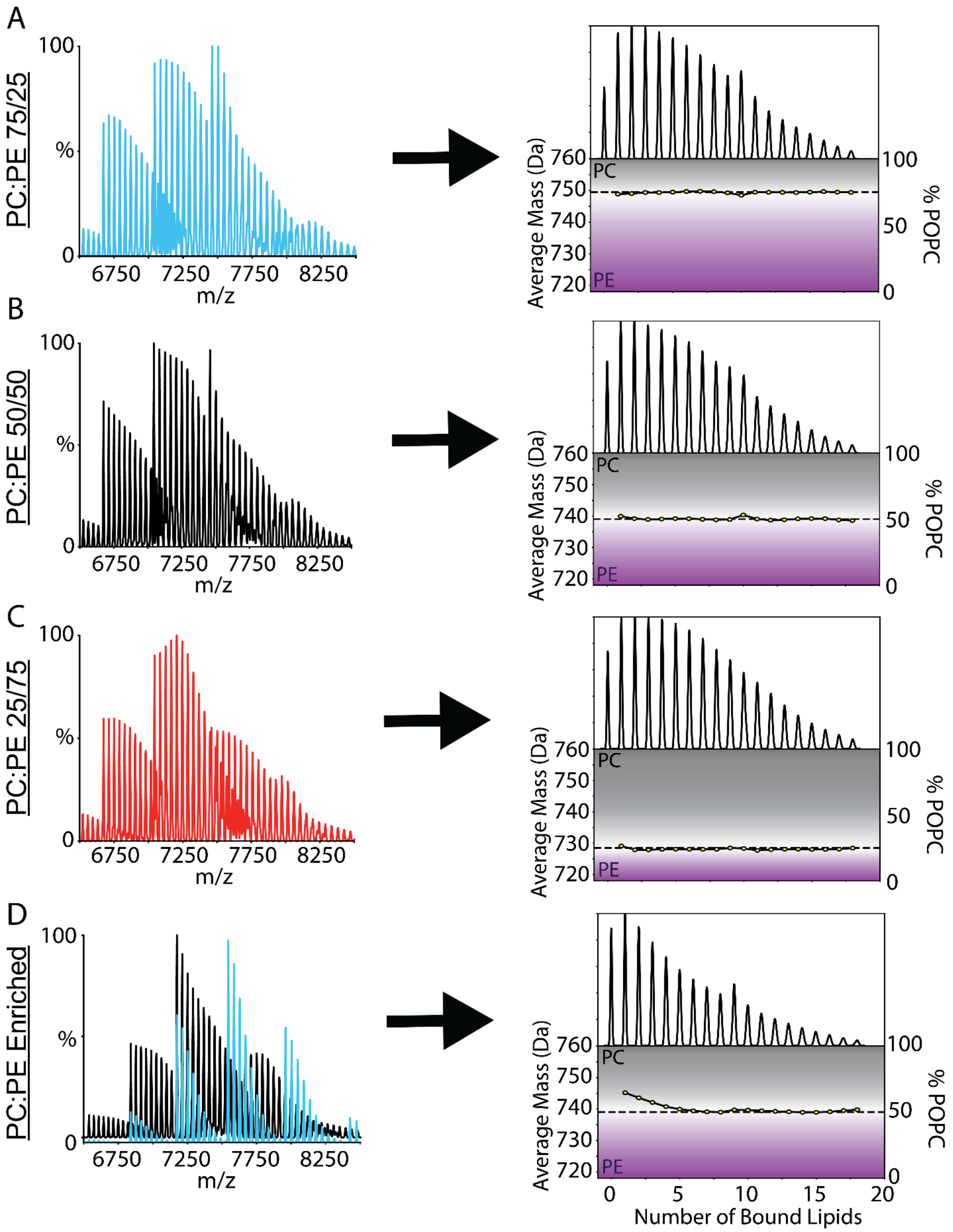

**Figure S-14**. Simulated mass spectra (A, B, C, D) of AmtB-lipid complexes ejected or extracted from 75/25 PC:PE (*blue*, A), 50/50 PC:PE (*black*, B), and 25/75 PC:PE (*red*, C) nanodiscs with no enrichment. A composite mass spectrum (D) was simulated by combining a mass spectrum corresponding to 75/25 PC:PE (*blue*) with lower numbers of bound lipids and a mass spectrum corresponding to 50/50 PC:PE (*black*) with higher numbers of bound lipids. The deconvolved spectra and corresponding average masses of bound lipids are indicated to the right. The expected average lipid masses are indicated by dashed lines.
